# Modern, archaeological, and paleontological DNA analysis of a human-harvested marine gastropod (*Strombus pugilis*) from Caribbean Panama

**DOI:** 10.1101/2020.08.26.269308

**Authors:** Alexis P. Sullivan, Stephanie Marciniak, Aaron O’Dea, Thomas A. Wake, George H. Perry

**Affiliations:** Departments of Biology, Pennsylvania State University, University Park, PA, USA; Departments of Anthropology, Pennsylvania State University, University Park, PA, USA; Smithsonian Tropical Research Institute, Panama City, Panama; Department of Anthropology and Costen Institute of Archaeology (CIOA), University of California, Los Angeles, CA, USA; Huck Institutes of the Life Sciences, Pennsylvania State University, University Park, PA, USA

**Author notes:** Addresses for correspondence.

## Abstract

Although protocols exist for the recovery of ancient DNA from land snail and marine bivalve shells, marine conch shells have yet to be studied from a paleogenomic perspective. We first present reference assemblies for both a 623.7 Mbp nuclear genome and a 15.4 kbp mitochondrial genome for *Strombus pugilis*, the West Indian fighting conch. We next detail a method to extract and sequence DNA from conch shells and apply it to conch from Bocas del Toro, Panama across three time periods: recently-eaten and discarded (n=3), Late Holocene (984-1258 BP) archaeological midden (n=5), and a mid-Holocene (5711-7187 BP) paleontological fossil coral reef (n=5). These results are compared to control DNA extracted from live-caught tissue and fresh shells (n=5). Using high-throughput sequencing, we were able to obtain *S. pugilis* nuclear sequence reads from shells across all age periods: up to 92.5 thousand filtered reads per sample in live-caught shell material, 4.57 thousand for modern discarded shells, 12.1 thousand reads for archaeological shells, and 114 reads in paleontological shells. We confirmed authenticity of the ancient DNA recovered from the archaeological and paleontological shells based on 5.7x higher average frequency of deamination-driven misincorporations and 15% shorter average read lengths compared to the modern shells. Reads also mapped to the *S. pugilis* mitochondrial genome for all but the paleontological shells, with consistent ratios of mitochondrial to nuclear mapped reads across sample types. Our methods can be applied to diverse archaeological sites to facilitate reconstructions of the long-term impacts of human behavior on mollusc evolutionary biology.

## INTRODUCTION

Human behavior has directly or indirectly affected non-human morphological evolution in many ways, well beyond plant and animal domestication (Sullivan, Bird, & Perry, 2017). One of the most well-studied of these mechanisms is that of non-human body size evolution due to size-selective human hunting and harvesting (C. T. Darimont, Fox, Bryan, & Reimchen, 2015; Chris T. Darimont et al., 2009). For example, intertidal mollusc exploitation by humans is well-documented throughout the archaeological record due to the trash heaps, or ‘middens’, of shells and other inedible materials deposited after processing (Erlandson & Rick, 2010). Routine, large-scale shellfish acquisition for sustenance has been recorded from ~120,000 years before present (BP) in South Africa (Jerardino, 2016), and the exploitation of this valuable resource continues to generate middens in certain areas of the world today (Bird & Bliege Bird, 1997; O’Dea, Shaffer, Doughty, Wake, & Rodriguez, 2014). While providing a record of human harvesting, these middens are also invaluable resources that can provide evidence of prey phenotypic change in response to these harvesting pressures through the ability to identify and quantify phyletic dwarfism over time (Avery et al., 2008; Erlandson et al., 2011; Klein & Steele, 2013; Stiner, Munro, Surovell, Tchernov, & Bar-Yosef, 1999).

Interest in obtaining DNA from molluscan shells has increased over the past few years due to their abundance in museum collections and recovery rate in the wild (Geist, Wunderlich, & Kuehn, 2008). By utilizing ancient DNA extraction and processing techniques, it is now possible to recover mitochondrial (Villanea, Parent, & Kemp, 2016) and even nuclear (Der Sarkissian et al., 2020, 2017) DNA from shell materials, even from those that are thousands of years old. This DNA could then be utilized for analyzing the genomes of these species to infer population history, biogeography, and more, even in non-human, non-model organisms (Coutellec, 2017; Perry, 2014). Once recovered from the shells, these temporal sequences of genetic records could potentially be used to confirm a genetic basis for the molluscan prey phenotypic change in response to human harvesting pressures and powerfully test adaptive hypotheses.

In 2014, O’Dea et al. (2014) published a study in which they analyzed West Indian fighting conch (*Strombus pugilis*) body sizes from modern, prehistoric midden (540-1260 BP; Wake et al., 2013), and paleontological reef (not human harvested; 7187-5711 BP; Fredston-Hermann, O’Dea, Rodriguez, Thompson, & Todd, 2013) sites in the Bocas del Toro archipelago in Caribbean Panama. Even though *S. pugilis* is smaller than its sympatric relative *Lobatus gigas* (queen conch), *S. pugilis* greatly outnumbers *L. gigas* and has remained an important component of the subsistence diet of the local people for millennia, as evidenced by its overwhelming presence in Bocas del Toro (Fredston-Hermann et al. 2013; Wake et al. 2013) and year-round harvesting and consumption today. Presently shells are either discarded in mounds near the harvesters’ houses or sold as souvenirs to tourists in gift shops (O’Dea et al., 2014).

By measuring the height, width, and lip thickness of *S. pugilis* shells from archaeological Sitio Drago (dated to AD 690-1410; Wake 2006; Wake et al. 2012, 2013), paleontological shells from a mid-Holocene fringing coral reef at Lennond (Fredston-Hermann et al., 2013; Lin et al., 2019; O’Dea et al., 2020), and modern sites spaced throughout the Bocas del Toro archipelago, O’Dea et al (2014) found that size at sexual maturity has decreased consecutively from the paleontological ‘pre-human’ period to the archaeological deposits to the present day. This size decrease is associated with a decline in edible meat weight by ~40% over the past 7,000 years (O’Dea et al., 2014).

For our present study, we revisited the sites and collections studied by O’Dea *et al* (2014) (see **Figure 1 A**) with the initial goal of assembling both a reference nuclear and mitochondrial genome for *S. pugilis* from freshly-preserved tissue. We then evaluated several DNA extraction techniques that were developed for mineralized substrates like ancient bone and bivalve shell materials (Der Sarkissian et al., 2017; Gamba et al., 2016, 2014; Kemp et al., 2007; Villanea et al., 2016; Yang, Eng, Waye, Dudar, & Saunders, 1998), and developed a modified method to extract modern and ancient DNA from the more robust recently-deceased and Holocene conch shells. We report our assessment of the mapping rates and DNA damage from modern live-caught, recently discarded, Late Holocene archaeological, and mid-Holocene paleontological shell materials.

**Figure 1:**
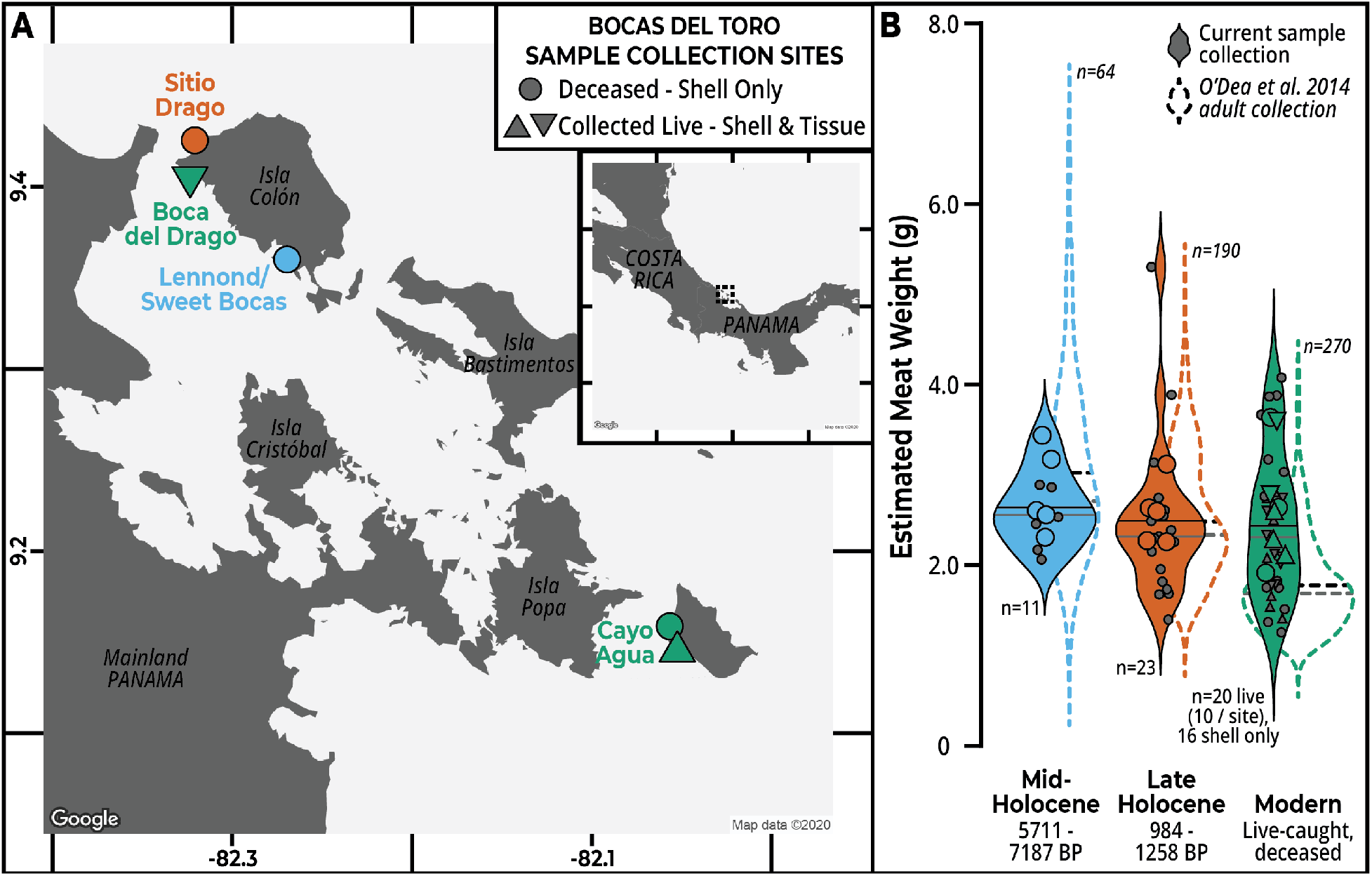
*Strombus pugilis* study populations. **A – *S. pugilis* specimen collection sites**. Live fighting conch individuals were collected from Boca del Drago and Cayo Agua 1 (green triangles), and shells from recently-eaten fighting conch were collected at Cayo Agua 2 (green circle). Late Holocene, archaeological, human-processed shells were collected from excavations at Sitio Drago (orange circle), and Mid-Holocene, pre-human shells from the exposed mid-Holocene fringing reef at the old town of Lennond, now called Sweet Bocas (blue circle). **B – Estimated meat weight distributions for adult *S. pugilis* specimens**. Height, width, and lip thickness measurements were collected for each shell, and calculations for estimated edible meat weight were calculated as described by O’Dea et al (2014). Filled violin plots represent distributions of specimens collected for this study, and unfilled plots depict adult shell distributions of shells presented in by O’Dea et al (2014; 144 from CA and 126 from Boca del Drago). The mean and median of each distribution are represented by the black and gray lines, respectively. Those individuals with larger points and colored interiors were sequenced for this study.

## MATERIALS AND METHODS

### Sample Collection Sites

We collected live *Strombus pugilis* individuals (n=10 each) for paired fresh tissue and shell subsamples from two of the contemporary sites sampled by O’Dea et al (2014): Cayo Agua (CA1, 9°08’46.6” N 82°03’8.9” W) and Boca del Drago (BD, 9°24’13.4” N 82°19’24.4” W; see **Figure 1 A**). Individual conchs were sited while snorkeling at the surface, then gathered from depths of approximately 4-6 meters. Sexually mature conchs were identified by an outer shell lip thickness of >1.8 mm (O’Dea et al., 2014); those with lip thicknesses less than or equal to 1.8 mm were released where they were caught. The live-caught mature individuals were transported in buckets to the Smithsonian Tropical Research Institute (STRI) Bocas del Toro Research Station (BDT) for subsampling and storage.

Recently discarded shells were collected from underneath a dock at CA1 (n=5) as well as from refuse piles at a second Cayo Agua site (CA2, 9°09’31.5” N 82°03’23.9” W; n=15), with the permission of the people living in each location. According to conversations with the harvesters, these conchs had been boiled inside their shells in water, the flesh removed for consumption, and the shells and any remaining attached tissue discarded into the domestic waste piles from which they were collected. The shells of these discarded individuals were rinsed with water and frozen whole in plastic zip bags at −20°C.

The Late Holocene archaeological and mid-Holocene paleontological shells were sourced from collections housed at STRI’s BDT and the Earl S. Tupper Research Center, respectively. The archaeological site “Sitio Drago” (see **Figure 1**) has been excavated in a series of 1×1 m units, and materials collected from them were sorted into two ceramic phases based on their depth in 10-cm levels: the Bisquit Ware phase (AD 1100–AD 1400, 0-40 cm) and the pre-Bisquit Ware phase (690–AD 1200, 40-150+ cm) (Wake 2006; Wake et al. 2013). We amassed all of the whole adult *S. pugilis* shells that were collected from units 60 and 61 (9°24’58.7” N 82°18’59.9” W; n=23 total, 9 from U60 and 14 from U61).

The paleontological *S. pugilis* remains (n=11 adult shells) were collected from a fossilized fringing reef excavated for construction purposes near the old town of Lennond (9°21’37.0” N 82°16’09.9” W; see **Figure 1**). Twenty four Uranium-Thorium and eight radiocarbon dates from 32 coral pieces date the reef at Lennond to the mid-Holocene (Mean = 6507, SD = 407, Min = 5711, Max = 7187 years BP; Fredston-Hermann et al., 2013; Lin et al., 2019; O’Dea et al., 2020; Dillon et al., in submission). This predates the earliest evidence of human interactions with marine species in the region which starts around around 4000 BP (Baldi, 2011), and confirms previous conclusions that the Caribbean slope of Panama was prehistorically not extensively populated until the Late Holocene (Griggs, 2005; O. F. Linares & White, 1980; Olga F. Linares, 1977; Piperno, Bush, & Colinvaux, 1990; Ranere & Cooke, 1991; Wake et al., 2013).

### Specimen Subsampling

The live *S. pugilis* individuals collected from each of the modern sites were kept in outdoor aquaria at the BDT Research Station prior to processing in the Station’s wet laboratory. The conchs were transferred to buckets, covered with seawater, and relaxed with 10 mL of a 2:1 solution of menthol oil and 95% ethanol (Sturm, Pearce, & Valdés, 2006). A set of forceps, a scalpel, and the dissection surface were sanitized with bleach (3-4% sodium hypochlorite and <1% sodium hydroxide), then washed with freshly prepared 70% ethanol to remove the bleach residue between samples. Once the conchs were relaxed (~3 hours), forceps were used to grasp the operculum and remove the conch from its shell. The operculum was removed from the main body of the conch prior to subsampling a 1-cm disk of foot muscle tissue. The tissue disks were stored individually submerged in RNAlater in 1.5-mL microcentrifuge tubes at room temperature.

All of the modern *S. pugilis* shells, from both the live-caught and recently discarded individuals, were rinsed with water, measured for height, width, and lip thickness (O’Dea et al., 2014; see **Figure 1 B, Supplementary Table 1**), then stored individually in plastic zip bags and frozen at −20°C. The archaeological and paleontological whole shells were also measured and stored in individual bags at room temperature. The shells were transported to the Pennsylvania State University for subsampling in an open-air facility. The subsampling surface and hood, forceps, and respective subsampling tool were cleaned with a 10% bleach solution followed by freshly prepared 70% ethanol between samples. The modern conch shells were rinsed with water to remove any residual frozen tissue or sand, and then the outer lip of each shell was sliced off in its entirety and stored in individual 50-mL falcon tubes at room temperature (see **Figure 2**).

**Figure 2:**
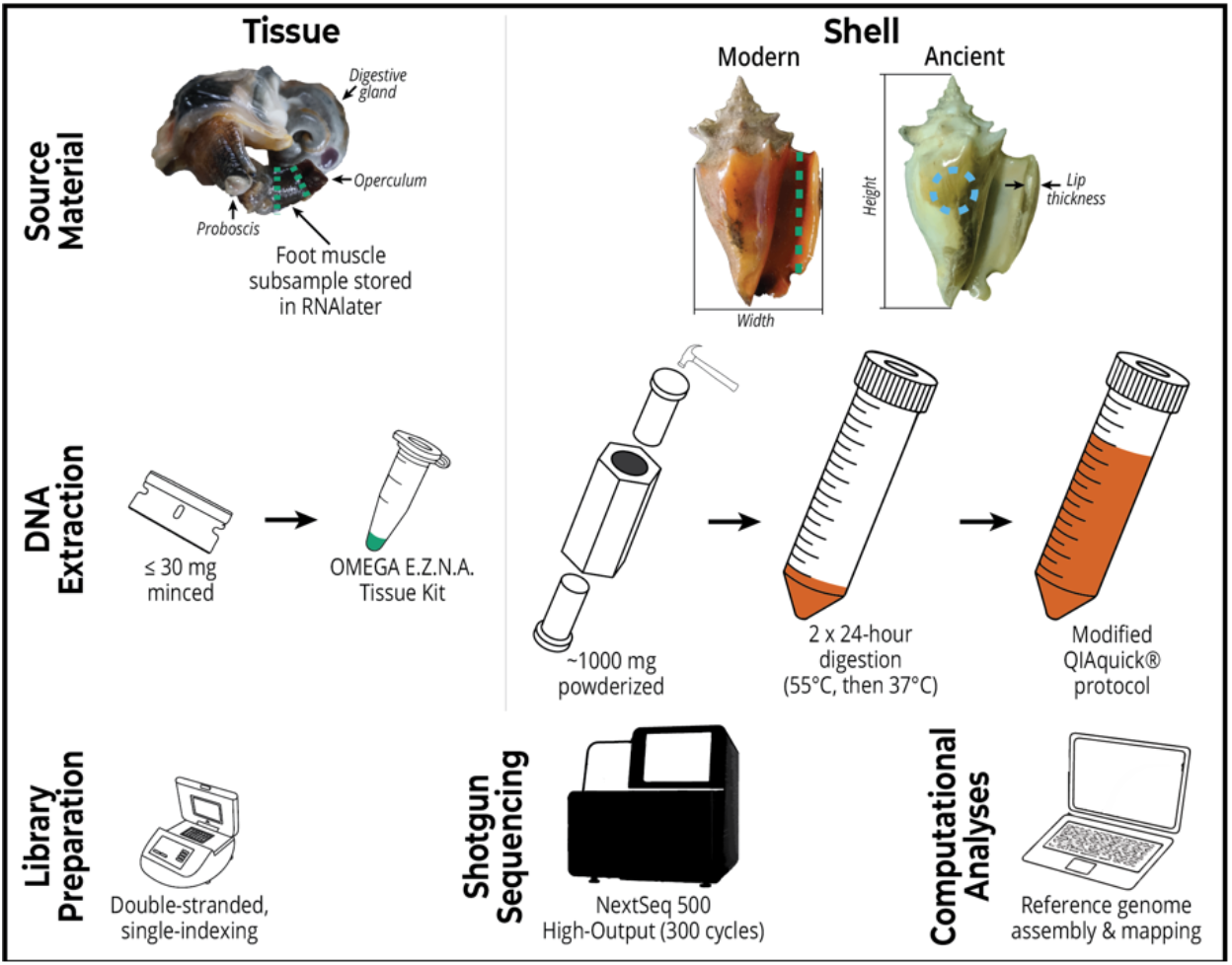
Protocol summary for *S. pugilis* DNA extraction, library preparation, sequencing, and data analysis.

The Late Holocene archaeological conch shells were also brushed and rinsed with water to remove any adhering sand and soil. All of the archaeological and paleontological shells were individually measured (see **Figure 1 B, Supplementary Table 1**) before a new sterilized ½-inch diamond-coated drill bit was used to remove a portion of the outer whorl of each conch shell (see **Figure 2**). These portions were stored in individual 15-mL falcon tubes at room temperature.

### Extraction Protocols

The subsampled preserved soft tissues and outer lip shell segments of the modern *S. pugilis* individuals were transferred to the Anthropological Genomics Laboratory for DNA extraction. The Holocene archaeological and paleontological subsamples were processed in a designated clean ancient DNA laboratory. Neither Penn State laboratory was exposed to molluscan material prior to this project, and extraction blank controls were incorporated in every step to monitor contamination.

We minced preserved *S. pugilis* foot muscle tissue (≤30 mg) for standard E.Z.N.A.Tissue Kit (OMEGA bio-tek) extractions (see **Supplementary Protocol 1** for full procedure). Trials were performed with the modern live-caught shells to test effectiveness of surface decontamination, extractions from several regions of the shell, amounts of starting material, extraction kits, and digestion time (Der Sarkissian et al., 2017; Gamba et al., 2016, 2014; Kemp et al., 2007; Villanea et al., 2016; Yang et al., 1998). Based on positive preliminary results, the following protocol was utilized for all of the subsampled shell materials in the respective laboratory spaces.

We prepared the *S. pugilis* shell fragments for extraction by removing the exterior surface of each shell with 220 grit sandpaper, rinsing throughout with deionized (DI) water, for decontamination. A customized stainless steel “shell smasher” was cleaned with detergent and DI water, disinfected with 10% bleach, rinsed with freshly prepared 70% ethanol, and dried before each new sample. The shell fragments were placed into the central chamber of the smasher and reduced to a powder, then ~1 g was transferred to a 50-mL falcon tube. Extraction buffer (0.5 M pH 8 EDTA, 0.5% sodium dodecyl sulfate, 0.25 mg/mL proteinase K) was prepared and warmed at 37°C until all precipitate had dissolved, then 4 mL was added to each sample. The samples were vortexed thoroughly then incubated at 55°C in a shaking heat block (≥ 750 rpm) for at least 24 hours.

After this first digestion, the samples were centrifuged at 1500 x *g* for 2 minutes, and the supernatant transferred to a 15-mL falcon tube without disturbing or transferring any of the insoluble pellet or remaining shell. The first digestion aliquot was stored at −20°C, and a second digestion was performed on the remaining shell material at 37°C for at least 24 hours. The samples were centrifuged at 2000 x *g* for 5 minutes, then the second digestion aliquot was combined with the first, without transferring the remaining shell pellet.

The combined aliquots were centrifuged at maximum speed (2520 x *g*) for 5 minutes, and the supernatant transferred into a 50-mL falcon tube. Five volumes of QIAquick PB Buffer were added to each sample, followed by 700 μL of sodium acetate (3 M, pH 5.5) and 15 μL of Pellet Paint. This extraction solution was filtered through QIAquick spin columns fitted with Zymo-Spin V extenders secured in a vacuum manifold. Once all of the extraction solution for each sample had passed through the column, the QIAquick spin columns were transferred to the reserved 2-mL Collection Tubes. The DNA was washed on the column with 750 μL of QIAquick PE buffer, then centrifuged at 12,800 x *g* for 1 minute. The filtrate was discarded, then the empty spin columns were centrifuged at 12,800 x *g* for 1 minute to dry the columns. We aliquoted the DNA with two 25-μL portions of 55°C-heated nuclease-free water. DNA concentrations for all samples were obtained with a Qubit 3.0 dsDNA High Sensitivity Kit, and the extractions stored at −20°C (see **Supplementary Protocol 1** for full procedure).

### Library Preparation and Sequencing

As with the DNA extractions, the modern samples were processed in the Anthropological Genomics Laboratory and the ancient samples in the designated clean ancient DNA laboratory. Libraries were prepared for each DNA extraction following a slightly modified version of the Meyer and Kircher 2010 protocol. 2000 ng of each tissue DNA sample was sheared for 48 seconds in a 115-μL suspension with a Covaris M220 ultrasonicator (peak incident power 50, duty factor 20%, 200 cycles per burst, temperature 20°C). The shell DNA, even for the modern samples, was not long enough to merit shearing since the initial step of library preparation involved a size-selection bead cleanup step. 50 μL of each sheared tissue DNA sample and 25 μL of each shell DNA sample, modern or ancient, was brought into library preparation. The volumes for the blunt-end repair, single-adapter ligation, adapter fill-in, and indexing PCR master mixes were kept the same for both the tissue and shell library preparations (see **Supplementary Protocol 2**). The *S. pugilis* DNA libraries were shotgun sequenced on a NextSeq500 High Output 300 cycles in two batches: one for the five randomly-selected tissue samples, and one for the 18 randomly-selected shell samples. One random tissue sample, Cayo Agua 2.3, was selected and pooled accordingly for high-coverage sequencing to generate the reference nuclear and mitochondrial assemblies.

### Reference Genome Assemblies

The Cayo Agua 2.3 reads for de novo reference assembly were trimmed with Trimmomatic (TruSeq2-PE adapters; Bolger, Lohse, & Usadel, 2014), and PCR duplicates were removed with filterPCRdupl.pl by Linnéa Smeds (https://github.com/linneas/condetri/blob/master/filterPCRdupl.pl). Kraken2 was used to taxonomically classify the trimmed reads, and, to avoid exogenous contamination, only those reads that were designated “unclassified”, “other sequences”, and “cellular organisms” were brought into assembly with soapdenovo v2.04 (kmer length 63; Wood, Lu, & Langmead, 2019). We removed all contigs smaller than 500 bp from the nuclear assembly prior to read mapping. We evaluated the quality of this nuclear assembly with QUAST, which determines assembly length, number of contigs, GC content, and contig N50, i.e. the minimum contig length that makes up half of the genome sequence (Gurevich, Saveliev, Vyahhi, & Tesler, 2013). The same Kraken2-classified Cayo Agua 2.3 reads were also used for norgal mitochondrial genome assembly, with the nuclear genome assembly provided to skip the initial de novo assembly generated by norgal (Al-Nakeeb, Petersen, & Sicheritz-Pontén, 2017). Both the nuclear and mitochondrial assemblies are available in SRA BioProject: PRJNA655996.

### Read Mapping

All raw sequence reads are available in SRA BioProject: PRJNA655996. The raw read sequence data for both our *S. pugilis* shell and tissue samples were assessed with FastQC (https://www.bioinformatics.babraham.ac.uk/projects/fastqc/) then trimmed with leehom, with the – ancientdna flag for all of the Late Holocene archaeological and mid-Holocene paleontological shells (Renaud, Stenzel, & Kelso, 2014). The tissue sequences were mapped to our nuclear and mitochondrial assemblies with BWA v0.7.16 *mem* (Li, 2013), while the modern and ancient shells were mapped to each assembly with BWA v0.7.16 *aln* since *aln* has a higher mapping rate for shorter reads (Li & Durbin, 2009). Once the reads for each sample were mapped, SAMtools v1.5 was used to convert the mapped SAM files to BAMs, sort the BAM files, remove duplicates, and filter the reads (minimum mapping quality of 30 and 30 base pair minimum length; Li et al., 2009). SAMtools v1.5 flagstat was used to count the number of alignments for every sample mapped to each assembly (Li et al., 2009).

### Damage Characterization

mapDamage v2.0.8-1 was used to characterize ancient shell DNA in terms of fragmentation patterns, nucleotide mis-incorporation, and fragment size distributions (see **Figure 3 B**; Jónsson, Ginolhac, Schubert, Johnson, & Orlando, 2013). Specifically, we quantified patterns of the rate of nucleotide (base) misincorporation, fragment length distributions, and strand fragmentation (due to depurination before sequence read starts). Deamination of cytosine to uracil due to hydrolytic damage preferentially occurs at read ends and causes miscoding lesions, specifically C>T (cytosine to thymine) misincorporations on the 5’ end and G>A (guanine to adenine) on the 3’ strand (Briggs et al., 2007; Dabney, Meyer, & Paabo, 2013). The elevated rates of nucleotide misincorporation in degraded/ancient material are a proxy of sequence validity, as modern DNA does not have this damage pattern (Hofreiter, 2001; Krause et al., 2010). Post-mortem, DNA is enzymatically degraded into smaller molecules leading to an excess of short DNA fragments relative to longer DNA fragments, which contributes to a characteristic distribution of reads for DNA sequences from ancient material (Molak & Ho, 2011; Paabo, 1989). Strand fragmentation in DNA reads manifests as an increase of purines (due to post-mortem depurination or the loss of adenine and guanine bases) before read starts (Briggs et al., 2007; Ginolhac, Rasmussen, Gilbert, Willerslev, & Orlando, 2011). The unique quality-filtered mapped read data for each shell category was combined with SAMtools v1.5 merge and filtered for duplicates for the total analysis (Li et al., 2009).

**Figure 3:**
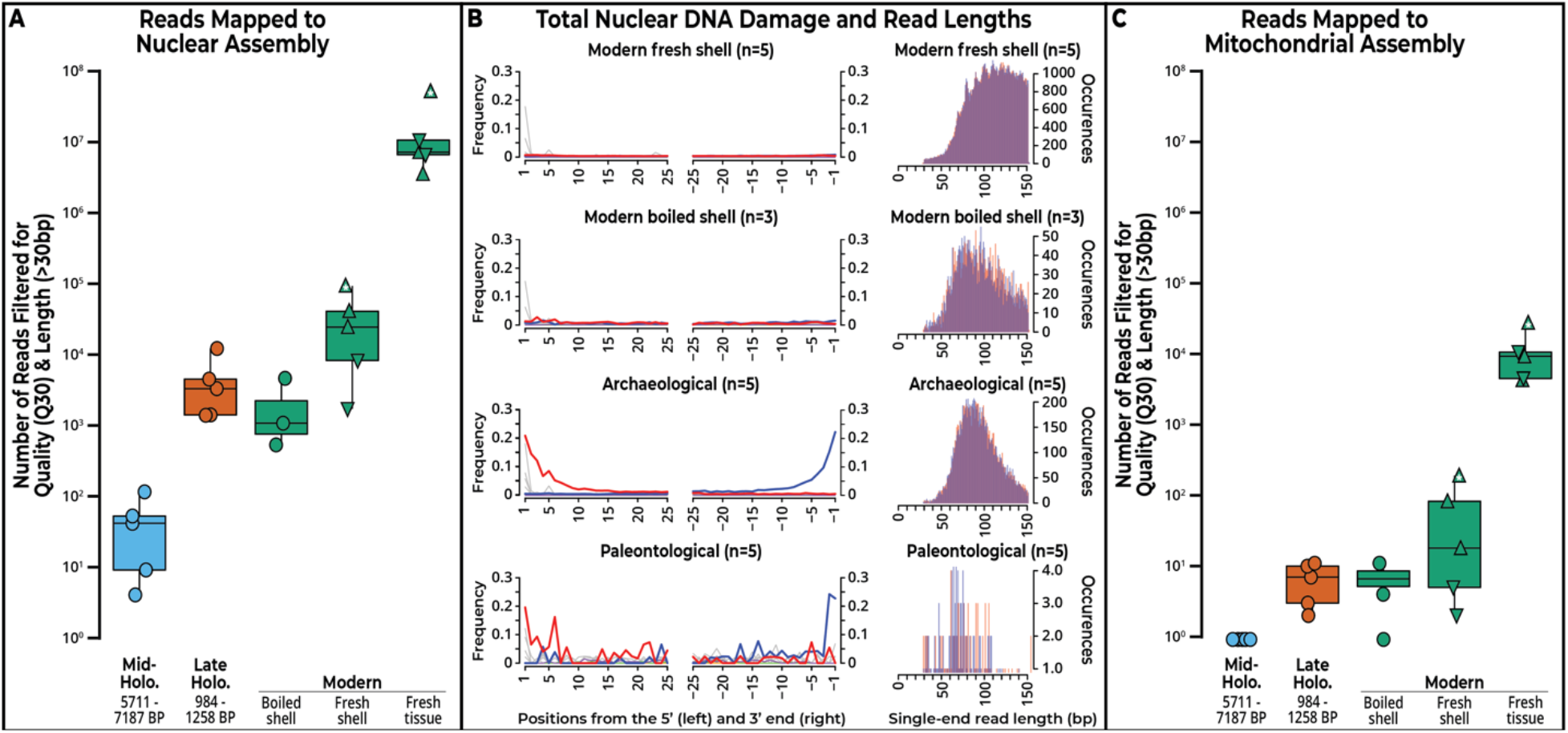
Recovery of *Strombus pugilis* nuclear and mitochondrial DNA from shells of varying ages. **A – Reads mapped to our *S. pugilis* nuclear assembly**. The reference nuclear genome was assembled from the conch individual marked with a white asterisk, Cayo Agua 2.3. *S. pugilis* fresh tissue and fresh shell samples were collected from the same individuals. Tissue samples were mapped to the reference with BWA *mem*, all shells were mapped with BWA *aln*. **B – Total nuclear DNA damage patterns and single-end read lengths for each group of *S. pugilis* shells**. The mapped bam files were merged for each age category of shells for mapDamage nucleotide mis-incorporation analysis. Color codes for the misincorporation plots: C-to-T substitutions in red, G-to-A in blue, all other substitutions in gray, soft-clipped bases in orange, deletions or insertions relative to the reference in green or purple, respectively. Color codes for the length plots: positive DNA strands in red, negative strands in blue. **C – Reads mapped to our *S. pugilis* mitochondrial assembly**. The reference mitochondrial genome was assembled from the conch individual marked with a white asterisk, Cayo Agua 2.3. *S. pugilis* fresh tissue and fresh shell samples were collected from the same individuals. Tissue samples were mapped to the reference with BWA *mem*, all shells were mapped with BWA *aln*.

## RESULTS

In this study we adapted existing DNA extraction protocols for use with the robust crystalline calcium matrix of conch shells. One of our primary motivations for doing so was to assess the amount and quality of DNA preserved in shell samples recovered from Late Holocene archaeological and mid-Holocene paleontological contexts. However, to be able to assess the effectiveness of our methods, we first needed to sequence and assemble both nuclear and mitochondrial reference genomes from a freshly preserved (modern) muscle tissue sample. Secondly, to assess the recovery of high-quality DNA from the mineralized shell matrix using our extraction protocol, we also generated sequencing data from DNA extracted from both modern conch muscle tissue and shell as baseline comparisons for the ancient DNA samples.

### Reference Genome Assemblies

Genomic data for *Strombus* species are limited, with no complete genome for *S. pugilis* available prior to our study. We used soapdenovo v2.04 to assemble a de novo reference nuclear genome sequence from the shotgun-sequenced reads of a randomly-selected *S. pugilis* individual (Cayo Agua 2.3). A total of 125,083,906 raw sequence reads were used to generate a nuclear assembly with a total length of 623,741,713 base pairs (bp), which is within the range of genome lengths expected of gastropods, 432.3 – 1,865.4 Mbp (Sun et al., 2019). Our *S. pugilis* nuclear assembly is composed of 697,168 contigs, with GC content at 44%. The contig N50 value for this assembly, or the minimum length of contigs that cover 50% of the genome sequence, is 908 bp (see **Supplementary Table 2** for full QUAST output). In the future, adding long-read sequencing data could help to increase the N50 value.

We also assembled a 15,409 bp mitochondrial genome for *S. pugilis* from the same Cayo Agua 2.3 high-coverage reads. The length of our reference-guided mitochondrial genome assembly is also within the range of those of other gastropods, 13,856 – 15,461 bp (Grande, Templado, & Zardoya, 2008; Márquez, Castro, & Alzate, 2016). The BLAST “best hit” for this assembly to a database of complete mtDNA and plastid genomes (Altschul, Gish, Miller, Myers, & Lipman, 1990; Camacho et al., 2009) was a 7,864 bp alignment to a published *Strombus* (now *Lobatus) gigas* mitochondrion (e-value 0, bit-score 6999; see **Supplementary Table 3** for full BLAST output; Márquez, Castro, and Alzate 2016).

### Developing a DNA Extraction Protocol for Modern and Ancient Conch Shells

Several DNA extraction techniques have been used for mineralized substrates like ancient bone, land snail, and marine bivalve shells (Der Sarkissian et al., 2017; Gamba et al., 2016, 2014; Kemp et al., 2007; Villanea et al., 2016; Yang et al., 1998). The protocol we describe here combines elements from each of these methods to address the difficulties in digesting robust conch shells. In brief, we initially digested 200 mg of freshly sanded and pulverized live-caught shell powder taken from the outer lip of the aperture (n=4) in 2 mL of SDS-based extraction buffer for 18 hours at 37°C and shaking at 750 rpm. SDS detergent is more effective than N-laurylsarcosyl in breaking down the mineralized matrix at room temperatures or higher. We added a second overnight shaking/heating step to digest more of the shell material: the samples first digested at 55°C, centrifuged to separate the extraction suspension and residual shell precipitate, their supernatant stored frozen at −20°C, and then a second overnight digest with fresh extraction buffer at 37°C. The initial fraction of supernatant was thawed at room temperature before being added back into the extraction solution. This addition of a two-day extraction period, plus Pellet Paint just before purification to help the DNA adhere to the spin column, yielded DNA for all samples.

Due to the potential for intra-site variability and choice of extraction parameters in impacting DNA yield, we used samples of shell from one live-caught *S. pugilis* individual to measure DNA yields from (i) multiple regions of the shell and (ii) QIAquick versus MinElute DNA purification from the digestion supernatant. To experimentally test multiple regions of the shell, along with the body whorl/aperture regions previously sampled, we ground shell from the protoconch (top-most whorl of the shell) and the siphonal canal since they are also areas where shell material is added during maturation. After following the digestion protocol previously mentioned with the addition of the third overnight digestion, half of the DNA extracts were purified with QIAquick spin columns and the other half with MinElute columns.

The MinElute spin-columns became visibly clogged and, likely as a consequence, purified less DNA from all of the shell regions than the QIAquick spin columns: average Qubit concentrations of 0.319±0.165 ng/μL and 0.757±0.204 ng/μL, respectively. The siphonal canal region of the shell yielded the least amount of DNA at 0.730 ng/μL, for QIAquick and 0.156 ng/μL for MinElute. The protoconch yielded concentrations of 0.974 ng/μL for QIAquick and 0.314 ng/μL for MinElute, though the protoconch proved more difficult to pulverize into fine-grained powder due to its rigid structure. Accordingly, using QIAquick silica spin columns and the easily accessible and most recently deposited outer lip shell material yielded the most DNA. The final adjustments included increasing the amount of starting material to 1.0 g of shell powder digested in 4-mL portions of the extraction buffer and using spin-column extenders during purification due to the buffer volume increase.

### Comparison Among Modern, Archaeological, and Paleontological Specimens

We mapped wholegenome shotgun sequence data from samples of freshly-preserved (modern) *S. pugilis* tissue and shells of varying ages (age range 0-7187 BP) to our nuclear reference assembly (see **Figure 3 A**; see **Supplementary Table 4** for filtered mapped read counts for all individuals). The muscle tissue sample group, including the one modern tissue sample we used to assemble our reference genome and four additional individuals, had an average of 15.9±20.1 million filtered reads map to the nuclear assembly out of 35.6±50.2 million total reads using the BWA long-read algorithm (v0.7.16, bwa-*mem*) (Li, 2013).

The sequencing libraries constructed from DNA extracted from modern and ancient conch shell samples (n=18) were pooled and sequenced with equimolar ratios, and then mapped to our nuclear reference genome using the BWA short-read algorithm bwa-*aln* v0.7.16 (Li & Durbin, 2009). On average, the fresh (modern) shell samples taken from the same individuals as the tissue samples had the highest number of unique quality-filtered reads map to the *S. pugilis* nuclear reference assembly (33.4±36.3 thousand of 2.30±0.89 million total reads sequenced). The recently eaten and discarded modern shell samples had fewer unique quality-filtered mapped reads than the archaeological shell samples with an average of 2.05±2.20 thousand (of 3.80±0.57 million total reads sequenced) and 4.51±4.42 thousand (of 3.56±1.06 million total reads sequenced), respectively. The proportions of reads that mapped to the nuclear reference assembly out of all sequenced reads from the discarded and archaeological shell samples are not significantly different (Welch two-sample t-test, p=0.2225). The shells from these groups were processed for consumption and then discarded either in soil or sea nearby, and their similar proportion of nuclear-mapped reads to total sequenced reads suggests long-term preservation of nuclear DNA in archaeological shells that is similar in quantity and quality to recently boiled shells. The paleontological shells had the fewest number of reads map to the nuclear assembly, with an average of just 44.0±44.2 (of 3.65±0.85 million total reads sequenced). The paleontological shells were likely exposed to much greater UV damage than any of the other shell groups, as evidenced by their bleached appearance.

We characterized patterns of post-mortem DNA damage to assess the likelihood of endogenous nuclear DNA authenticity for each of our groups of *S. pugilis* shells (**Figure 3 B**; see **Supplementary Figure 1** for individual shell results). As expected, the nuclear DNA reads for both of the modern groups of conch shells do not exhibit post-mortem DNA fragmentation patterns, though the fragment length distribution of trimmed single-end reads from the recently boiled and discarded shells are slightly shorter in length (average 96±27 bp) than those of the live-caught individuals (average 109±26 bp; **Figure 3 B**), which could be a potential impact from people boiling these shells. The authenticity of ancient DNA recovered from the archaeological and paleontological shells is confirmed based on the presence of typical molecular signatures of post-mortem DNA damage. Specifically, the sequence reads obtained from the ancient samples have shorter fragment lengths relative to those from the modern shells (average 97±28 bp for the archaeological shells, 76±28 bp for the paleontological; **Figure 3 B**).

The ancient samples also exhibit higher rates of C>T nucleotide misincorporations at 5’ ends (G>A at the 3’ ends) due to post-mortem cytosine-driven deamination than the modern samples, resulting in the classic aDNA “smile” fragmentation pattern: average frequency of 0.005 for live shells, 0.010 for boiled, 0.041 for archaeological, and 0.42 for paleontological (**Figure 3 B**). Comparably, the rate of cytosine deamination in double-stranded regions (δd) is 0.0164±0.0013 for archaeological and 0.0174±0.0083 for paleontological, with lower rates in live and boiled shells (0.0003±0.0003 and 0.0034±0.0023, respectively). The more jagged “smile” fragmentation pattern and size distribution of the paleontological shells is attributed to the fewer high-quality mapped reads compared to the archaeological shells, potentially due to environmental/taphonomic conditions (e.g., bleaching by the sun in the exposed reef).

We also mapped the tissue and shell sequence data to our mitochondrial reference assembly (see **Figure 3 C**; see **Supplementary Table 4** for filtered mapped read counts for all individuals). The modern tissue samples mapped with BWA *mem* averaged 11.5±9.7 thousand unique quality-filtered mapped reads, and the shell samples from those same modern individuals mapped with BWA *aln* averaged 60±81 reads. When both the live-caught tissue and shell samples were mapped with the same program, BWA *mem*, the proportions of filtered mitochondrial reads to total reads were similar across both sample sources (0.071% mitochondrial reads out of total filtered mapped reads for the tissue samples, 0.074% for the shell samples; Fisher’s exact test, p=0.3691).

The average proportions of filtered BWA *aln*-mapped mitochondrial reads relative to mitochondrial + nuclear mapped reads in the boiled and discarded modern shells (5.0±5.6) and archaeological shells (6.6±4.0) are similar to that observed for live-caught shell material (Fisher’s exact test; p=0.2247 and 0.3087, respectively). No filtered reads mapped to the mitochondrial genome for the paleontological shells, but this results still meets expectations given the ratio of nuclear:mtDNA mapped reads for modern shells (Fisher’s exact test; p=1), given the relatively small number of nuclear-mapped reads for the paleontological specimens.

## DISCUSSION

Genomics is an increasingly accessible tool for studying phylogenetics, trait evolution, adaptation, and population dynamics, and sequences for non-model organisms have been increasingly available over the past decade (Ellegren, 2014). Molluscan DNA in particular is receiving attention given the fact that molluscs are an economically important marine resource, and are important for ecological, evolutionary, zooarchaeological, and even mechanical studies because of their shells (Ferreira et al., 2020; Gomes-dos-Santos, Lopes-Lima, Castro, & Froufe, 2020). Molluscan shells, especially structurally robust conch shells, are a reservoir for both nuclear and mitochondrial DNA; their abundance in museum collections provides an opportunity to explore mollusc evolutionary change over time, including direct and indirect responses to human behavior. This study represents the first attempt to harness DNA from tropical marine shells.

Our results demonstrate that authentic endogenous DNA can be extracted from *Strombus pugilis* shells, including specimens up to ~7,000 years old. DNA extraction was most successful (in terms of the number of unique mapped reads) from modern (fresh and boiled) and archaeological conchs, while paleontological conchs (5711-7187 BP) yielded the lowest amount of DNA, potentially due to postmortem taphonomic conditions of UV exposure at the dry reef site. Macroscopically, the outer lip aperture of the mid-Holocene paleontological shells exhibited color bleaching and more brittle textures compared to the previously buried Late Holocene archaeological shells, as such we recommend researchers evaluate the integrity of this region when selecting samples for ancient DNA analysis. Our conch shell extraction protocol would be compatible with enrichment methods applications (e.g., hybridization-based DNA capture), which would further enhance the efficiency of nuclear and mitochondrial DNA reconstruction from archaeological and paleontological conch shells.

Our results confirm the long-term preservation of conch DNA in shell. This finding is consistent with those of other studies with molluscan shells dating to similar time periods (Der Sarkissian et al., 2017). With the ability to extract and sequence high-quality DNA from both modern and ancient conch shells, there is exciting potential in expanding the temporal range of evolutionary studies using these large marine snails across a range of contexts, including archaeological middens and pre-human sites. For our own research, we hope to identify genomic regions that correlate to the decreased body size phenotype via a genome-wide association study, use population genetics to test the hypothesis of natural selection on alleles associated with that phenotype, and incorporate ancient DNA for direct evidence of allele frequency change over time.

There is further potential for future studies on *S. pugilis* utilizing the methods presented here. *S. pugilis* ranges from the Northern Caribbean to Brazil and has experienced vastly different degrees of intensity and duration of human harvesting, as documented in archaeological shell deposits and historical documents across this range. In some regions this conch species was commonly consumed in the past but is now rarely eaten. In other regions *S. pugilis* remains a popular and economically important delicacy or is frequently harvested for its shell, which is frequently sold as a souvenir for tourists. In both cases it is highly likely that harvesters focus on the largest individuals. Unpublished data show that size at maturity varies considerably across these gradients of human selection. Although further study is needed, evidence exists suggesting that when harvesting pressure is removed from a population, *S. pugilis* has larger body sizes compared to adjacent populations that continue to be harvested (O’Dea et al., 2014). Paleo- and modern genomics studies of widely harvested animals such as *S. pugilis* could therefore represent an important tool to better understand the impacts of human-selective harvesting, and could guide management approaches to slow, or even reverse, deleterious effects of human-induced evolution in wild populations. These general methods can also be applied to fossil and archaeological material more broadly to facilitate new explorations into ancient metagenomic evolutionary biology of carbonate-bound DNA.

## ACKNOWLEDGMENTS

We thank Ashley Sharpe, Nicole Smith-Guzman, Suzette Flantua, Jarrod Scott, Matthieu Leray, and Wilmer Elvir for their assistance in live conch collection, and Brigida de Gracia for assisting with the planning and coordination at the STRI Naos Marine and Molecular Laboratories. We also thank the Bocas del Toro Research Station team, especially Plinio Góndola and Urania González. Thank you, Marian Shaffer and Felix Rodriguez, for helping to collect the paleontological conch, as well as Marco and Fausto Alvarez, Annick Belanger, and the staff at Sweet Bocas who kindly gave logistical support. Permits to collect were provided by three Panamanian ministries; Ministerio de Comercio e Industria (MICI), MiAmbiente, and El Ministerio de Cultura de Panamá (formerly Instituto Nacional de Cultura, INAC). We thank the Dirección Nacional del Patrimonio Histórico, Instituto Nacional de Cultura, Republica de Panamá for issuing Resolución 153-14, permitting the excavation of units 60 and 61 at Sitio Drago and the Serracín family for access to the site – especially Juany and Willy Serracín and Ana Serracín de Shaffer. We also thank Daniel Schussheim for machining the shell smasher, the NYU Langone Genome Technology Center for sequencing our libraries, and Christina Bergey, Kathleen Grogan, and the other members of the Perry Lab for their discussion and analytical advice. Components of this work were supported by the National Science Foundation Graduate Research Fellowship program (DGE-1255832, to A.P.S.), a grant from the National Science Foundation (BCS-1554834 to G.H.P.), the Smithsonian Tropical Research Institute Short-Term Fellowship Program (to A.P.S.), and the SNI program from the *Secretaría Nacional de Ciencia y Tecnología e Innovación*, Panamá (to A.O.).

## AUTHOR CONTRIBUTIONS

A.P.S, A.O., and G.H.P. conceived of and designed the study. A.P.S., A.O., and T.A.W. collected live *S. pugilis*, T.A.W. amassed the archaeological collection, and A.O. the paleontological collection. A.P.S. carried out the wet lab and computational work, with training, consultation, and input from S.M. A.P.S. drafted the manuscript, with contributions from all authors. All authors gave final approval for publication and agree to be held accountable for the work presented here.

## DATA, CODE, AND MATERIALS

Raw sequence reads and reference genomes are available at NCBI SRA BioProject: PRJNA655996. All code developed and used for this research work are stored in GitHub (https://github.com/AlexisPSullivan/Conch) and have been archived within the Zenodo repository (doi: 10.5281/zenodo.3993744).

## SUPPLEMENTARY MATERIALS

**Supplemental Database:** Poster (https://scholarsphere.psu.edu/concern/generic_works/41n79h518r)

**Supplemental Protocol 1**: Checklist-formatted protocol for DNA extraction from *Strombus pugilis* shells (https://scholarsphere.psu.edu/concern/generic_works/41n79h518r)

**Supplemental Protocol 2**: Checklist-formatted protocol for our modified library preparation protocol for single-indexed, whole-genome shotgun sequencing (https://scholarsphere.psu.edu/concern/generic_works/41n79h518r)

**Supplementary Table 1**: Morphometric data for all collected *Strombus pugilis* specimens

**Supplementary Table 2**: *S. pugilis* nuclear reference assembly QUAST metrics

**Supplementary Table 3**: *S. pugilis* mitochondrial reference assembly norgal BLAST metrics

**Supplementary Table 4**: Filtered mapped read count for all sequenced *S. pugilis* specimens

**Supplementary Figure 1**: Nuclear DNA damage and read length results for individual shell samples

