## Supplementary Materials for "Modern, archaeological, and paleontological DNA analysis of a human-harvested marine gastropod (*Strombus pugilis*) from Caribbean Panama"

**Supplemental Database:** Poster (https://scholarsphere.psu.edu/concern/generic_works/41n79h518r)

**Supplemental Protocol 1**: Checklist-formatted protocol for DNA extraction from *Strombus pugilis* shells (https://scholarsphere.psu.edu/concern/generic_works/41n79h518r)

**Supplemental Protocol 2**: Checklist-formatted protocol for our modified library preparation protocol for single-indexed, whole-genome shotgun sequencing (https://scholarsphere.psu.edu/concern/generic_works/41n79h518r)

**Supplementary Table 1**: Morphometric data for all collected *Strombus pugilis* specimens

**Supplementary Table 2**: *S. pugilis* nuclear reference assembly QUAST metrics

**Supplementary Table 3**: *S. pugilis* mitochondrial reference assembly norgal BLAST metrics

**Supplementary Table 4**: Filtered mapped read count for all sequenced *S. pugilis* specimens

**Supplementary Figure 1**: Nuclear DNA damage and read length results for individual shell samples

**Supplementary Table 1**: Morphometric data for all collected *Strombus pugilis* specimens

| *Individual* | *Site Age* | *Site Name* | *Length (mm)* | *Width (mm)* | *Lip Thickness (mm)* | *Estimated Meat Weight (g)* | *Sex* | *Shell Condition* |
|---|---|---|---|---|---|---|---|---|
| ModSite-1-1 | Modern | ModSite | 77.4 | 52.05 | 4.4 | 3.657219182 | Female | |
| ModSite-1-2 | Modern | ModSite | 83.14 | 52.44 | 4.32 | 4.693077546 | Female | |
| ModSite-1-3 | Modern | ModSite | 80.49 | 56.58 | 4.94 | 4.191952558 | Male | |
| ModSite-1-4 | Modern | ModSite | 75.92 | 50.8 | 3.42 | 3.419178006 | Female | |
| ModSite-1-5 | Modern | ModSite | 69.62 | 42.64 | 3.11 | 2.527963447 | Female | |
| ModSite-1-6 | Modern | ModSite | 78.41 | 53.15 | 4.02 | 3.826298858 | Male | |
| ModSite-1-7 | Modern | ModSite | 78.74 | 53.37 | 5.23 | 3.882730085 | Male | |
| ModSite-1-8 | Modern | ModSite | 85.21 | 54.58 | 4.77 | 5.113168569 | Female | |
| ModSite-1-9 | Modern | ModSite | 73.71 | 47.04 | 4.43 | 3.084587323 | Male | |
| ModSite-1-10 | Modern | ModSite | 82.79 | 53.33 | 4.4 | 4.624565114 | Female | |
| CayoAgua-2-1 | Modern | CayoAgua1 | 71.16 | 40.74 | 4.68 | 2.728314837 | Female | |
| CayoAgua-2-2 | Modern | CayoAgua1 | 69.31 | 43.2 | 5.27 | 2.488940436 | Male | |
| CayoAgua-2-3 | Modern | CayoAgua1 | 70.11 | 40.78 | 4.8 | 2.590532024 | Male | |
| CayoAgua-2-4 | Modern | CayoAgua1 | 61.55 | 36.97 | 4.59 | 1.645314813 | Male | |
| CayoAgua-2-5 | Modern | CayoAgua1 | 66.06 | 43.49 | 3.85 | 2.105260552 | Female | |
| CayoAgua-2-6 | Modern | CayoAgua1 | 67.57 | 44.05 | 5.48 | 2.277834473 | Male | |
| CayoAgua-2-7 | Modern | CayoAgua1 | 60.26 | 34.89 | 3.13 | 1.528204724 | Male | |
| CayoAgua-2-8 | Modern | CayoAgua1 | 58.72 | 33.66 | 3.38 | 1.396330291 | Male | |
| CayoAgua-2-9 | Modern | CayoAgua1 | 65.72 | 41.86 | 4.71 | 2.067729272 | Male | |
| CayoAgua-2-10 | Modern | CayoAgua1 | 71.64 | 47.88 | 3.99 | 2.793009044 | Female | |
| BocaDrago-3-1 | Modern | BocaDrago | 70.28 | 44.02 | 4.13 | 2.612495135 | Female | |
| BocaDrago-3-2 | Modern | BocaDrago | 68.12 | 40.64 | 3.1 | 2.343124648 | Female | |
| BocaDrago-3-3 | Modern | BocaDrago | 71.62 | 47.72 | 4.07 | 2.790291833 | Male | |
| BocaDrago-3-4 | Modern | BocaDrago | 63.14 | 39.1 | 4.44 | 1.798298398 | Male | |
| BocaDrago-3-5 | Modern | BocaDrago | 66.31 | 42.71 | 3.55 | 2.133165206 | Female | |
| BocaDrago-3-6 | Modern | BocaDrago | 66.08 | 41.58 | 3.33 | 2.107483289 | Male | |
| BocaDrago-3-7 | Modern | BocaDrago | 71.37 | 43.46 | 4.72 | 2.756485542 | Female | |
| BocaDrago-3-8 | Modern | BocaDrago | 63.07 | 39.26 | 3.58 | 1.791358005 | Male | |
| BocaDrago-3-9 | Modern | BocaDrago | 65.3 | 43.29 | 5.17 | 2.02202886 | Male | |
| BocaDrago-3-10 | Modern | BocaDrago | 77.08 | 49.21 | 4.94 | 3.604780184 | Female | |
| SitioDrago-U61-0_10-5223 | Archaeological | SitioDrago-U61 | 70.23 | 44.92 | 5.61 | 2.606021666 | NA | top dull |
| SitioDrago-U61-10_20-5305A | Archaeological | SitioDrago-U61 | 63.32 | 42.91 | 3.62 | 1.816233153 | NA | |
| SitioDrago-U61-10_20-5305B | Archaeological | SitioDrago-U61 | 62.46 | 39.2 | 3.4 | 1.731683432 | NA | bottom dull |
| SitioDrago-U61-20_30 | Archaeological | SitioDrago-U61 | 67.41 | 43.9 | 4.21 | 2.25908725 | NA | |
| SitioDrago-U61-50_60-5454 | Archaeological | SitioDrago-U61 | 73.93 | 47.2 | 3.98 | 3.116800334 | NA | |
| SitioDrago-U61-70_80 | Archaeological | SitioDrago-U61 | 78.75 | 47.45 | 4.61 | 3.88444933 | NA | |
| SitioDrago-U61-80_90-5596A | Archaeological | SitioDrago-U61 | 67.51 | 41.44 | 3.41 | 2.270791315 | NA | |
| SitioDrago-U61-80_90-5596B | Archaeological | SitioDrago-U61 | 66.46 | 40.52 | 4.65 | 2.150034034 | NA | |
| SitioDrago-U61-80_90-5596C | Archaeological | SitioDrago-U61 | 69.3 | 45.01 | 4.71 | 2.487688828 | NA | |
| SitioDrago-U61-90_100A | Archaeological | SitioDrago-U61 | 67.38 | 46.09 | 4.98 | 2.255584436 | NA | |
| SitioDrago-U61-90_100B | Archaeological | SitioDrago-U61 | 61.88 | 43.9 | 3.77 | 1.676271508 | NA | |
| SitioDrago-U61-100_110 | Archaeological | SitioDrago-U61 | 69.76 | 45.64 | 4.66 | 2.545728949 | NA | |
| SitioDrago-U61-110_120-5651A | Archaeological | SitioDrago-U61 | 71.28 | 43.14 | 4.01 | 2.744387111 | NA | |
| SitioDrago-U61-110_120-5651B | Archaeological | SitioDrago-U61 | 68.48 | 48.21 | 4.38 | 2.386575832 | NA | |
| SitioDrago-U60-10_20 | Archaeological | SitioDrago-U60 | 70.15 | 48.52 | 4.66 | 2.595687922 | NA | |
| SitioDrago-U60-20_30-5310 | Archaeological | SitioDrago-U60 | 64.67 | 44.68 | 4.13 | 1.954835302 | NA | |
| SitioDrago-U60-30_40A | Archaeological | SitioDrago-U60 | 74.06 | 49.14 | 4.82 | 3.135947657 | NA | |
| SitioDrago-U60-30_40B | Archaeological | SitioDrago-U60 | 58.72 | 39.61 | 3.75 | 1.396330291 | NA | |
| SitioDrago-U60-40_50 | Archaeological | SitioDrago-U60 | 86.09 | 55.76 | 3.51 | 5.299624992 | NA | |
| SitioDrago-U60-70_80-5530A | Archaeological | SitioDrago-U60 | 67.89 | 44.07 | 2.84 | 2.315661361 | NA | |
| SitioDrago-U60-70_80-5530B | Archaeological | SitioDrago-U60 | 67.72 | 43.08 | 4.1 | 2.295510511 | NA | |
| SitioDrago-U60-90_100-5564 | Archaeological | SitioDrago-U60 | 61.96 | 43.03 | 4.51 | 1.683838255 | NA | |
| SitioDrago-U60-110_120-5659 | Archaeological | SitioDrago-U60 | 70.44 | 46.6 | 4.72 | 2.6332873 | NA | |
| Lennond-SweetBocas-APS-MS-17-2-78 | Paleontological | Lennond-SweetBocas | 70.21 | 43.01 | 4.98 | 2.603435485 | NA | |
| Lennond-SweetBocas-APS17-MS-F-2-217 | Paleontological | Lennond-SweetBocas | 72.14 | 44.76 | 3.59 | 2.861554469 | NA | |
| Lennond-SweetBocas-APS17-MS-F-5-157 | Paleontological | Lennond-SweetBocas | 74.3 | 46.59 | 4.22 | 3.171516716 | NA | |
| Lennond-SweetBocas-APS17-MS-F-5-166 | Paleontological | Lennond-SweetBocas | 65.67 | 41.78 | 4.54 | 2.062250506 | NA | |
| Lennond-SweetBocas-APS17-MS-F-7-128 | Paleontological | Lennond-SweetBocas | 76.03 | 47.55 | 4.36 | 3.436478862 | NA | |
| Lennond-SweetBocas-APS17-MS-F-10-112 | Paleontological | Lennond-SweetBocas | 72.33 | 49.63 | 4.05 | 2.887913423 | NA | |
| Lennond-SweetBocas-APS17-MS-F-10-110 | Paleontological | Lennond-SweetBocas | 69.84 | 41.65 | 4.11 | 2.555920541 | NA | |
| Lennond-SweetBocas-APS17-MS-F-15-1 | Paleontological | Lennond-SweetBocas | 69.66 | 45.53 | 4.1 | 2.533030253 | NA | |
| Lennond-SweetBocas-APS17-MS-F-15-2 | Paleontological | Lennond-SweetBocas | 66.62 | 45.31 | 3.56 | 2.168132075 | NA | |
| Lennond-SweetBocas-APS17-MS-F-15-3 | Paleontological | Lennond-SweetBocas | 69.04 | 43.27 | 3.16 | 2.455304323 | NA | |
| Lennond-SweetBocas-APS17-MS-F-15-4 | Paleontological | Lennond-SweetBocas | 67.82 | 44.94 | 3.55 | 2.30734874 | NA | |
| CayoAgua-Boil1 | Modern | CayoAgua2 | 70.48 | 43.71 | 4.38 | 2.638503723 | NA | |
| CayoAgua-Boil2 | Modern | CayoAgua2 | 77.26 | 46.96 | 4.21 | 3.634210656 | NA | |
| CayoAgua-Boil3 | Modern | CayoAgua2 | 64.28 | 42.97 | 5.19 | 1.914046464 | NA | |
| CayoAgua-Boil4 | Modern | CayoAgua2 | 77.44 | 46.63 | 2.31 | 3.66381208 | NA | |
| CayoAgua-Boil5 | Modern | CayoAgua2 | 79.84 | 44.58 | 4.21 | 4.075123375 | NA | |
| CayoAgua-Boil6 | Modern | CayoAgua2 | 71.4 | 41.43 | 4.78 | 2.76052679 | NA | |
| CayoAgua-Boil7 | Modern | CayoAgua2 | 74.28 | 44.42 | 3.29 | 3.168541694 | NA | |
| CayoAgua-Boil8 | Modern | CayoAgua2 | 62.65 | 39.82 | 4.02 | 1.750116134 | NA | |
| CayoAgua-Boil9 | Modern | CayoAgua2 | 78.71 | 48.11 | 4.67 | 3.877575606 | NA | |
| CayoAgua-Boil10 | Modern | CayoAgua2 | 78.64 | 49.61 | 4.33 | 3.865567469 | NA | |
| CayoAgua-Boil11 | Modern | CayoAgua2 | 73.34 | 43.52 | 3.43 | 3.030947382 | NA | top/bottom dull |
| CayoAgua-Dock1 | Modern | CayoAgua1 | 56.94 | 35.46 | 4.86 | 1.254253366 | NA | bottom dull |
| CayoAgua-Dock2 | Modern | CayoAgua1 | 63.41 | 42.34 | 4.64 | 1.825248195 | NA | top/bottom dull |
| CayoAgua-Dock3 | Modern | CayoAgua1 | 60.07 | 39 | 4.11 | 1.511473388 | NA | top/bottom dull, lip broken |
| CayoAgua-Dock4 | Modern | CayoAgua1 | 58.4 | 35.49 | 4.4 | 1.369983022 | NA | top dull |
| CayoAgua-Dock5 | Modern | CayoAgua1 | 62.67 | 39.4 | 4.63 | 1.752064523 | NA | top dull |

**Supplementary Table 2**: *S. pugilis* nuclear reference assembly QUAST metrics

| *Assembly* | *Cayo_Agua_2-3_S1_kraken4* | *Cayo_Agua_2-3_S1_kraken4_broken* |
|---|---|---|
| *# contigs (>= 0 bp)* | 697,168 | - |
| *# contigs (>= 1000 bp)* | 180,494 | 80,727 |
| *# contigs (>= 5000 bp)* | 490 | 129 |
| *# contigs (>= 10000 bp)* | 7 | 2 |
| *# contigs (>= 25000 bp)* | 0 | 0 |
| *# contigs (>= 50000 bp)* | 0 | 0 |
| *Total length (>= 0 bp)* | 624,646,899 | - |
| *Total length (>= 1000 bp)* | 268,783,557 | 115,504,023 |
| *Total length (>= 5000 bp)* | 2,981,588 | 782,733 |
| *Total length (>= 10000 bp)* | 75,057 | 20,895 |
| *Total length (>= 25000 bp)* | 0 | 0 |
| *Total length (>= 50000 bp)* | 0 | 0 |
| *# contigs* | 695,354 | 478,846 |
| *Largest contig* | 12,028 | 10,766 |
| *Total length* | 623,741,713 | 379,042,087 |
| *GC (%)* | 44 | 44 |
| *N50* | 908 | 771 |
| *N75* | 673 | 606 |
| *L50* | 225,771 | 165,931 |
| *L75* | 427,335 | 305,789 |
| *# N's per 100 kbp* | 14,883 | 0 |

#### **Supplementary Table 3**: *S. pugilis* mitochondrial reference assembly norgal BLAST metrics

| *Type* | *Scaffold:Scaffold-length* | *Identity* | *Alignment-length* | *Ref.Length* | *E-value* | *Bit-score* | *Best-hit reference* |
|---|---|---|---|---|---|---|---|
| m | scaffold_0:15409 bp | 82.93 | 7864 | 15461 | 0 | 6999 | Strombus gigas isolate Sg300-UNAL-SAA mitochondrion, complete genome |
| p | scaffold_79678:1116 bp | 87.05 | 139 | 156073 | 6.00E-35 | 152 | Nicotiana otophora chloroplast, complete genome |
| m | scaffold_27902:946 bp | 88.50 | 113 | 17572 | 2.00E-29 | 134 | Anomaloglossus baeobatrachus voucher AF2590 mitochondrion, complete genome |
| p | scaffold_20072:772 bp | 94.52 | 73 | 153429 | 2.00E-23 | 113 | Bryopsis hypnoides chloroplast, complete genome |
| m | scaffold_51871:610 bp | 95.08 | 61 | 16000 | 1.00E-18 | 97.1 | Fergusonina taylori mitochondrion, complete genome |
| m | scaffold_27962:714 bp | 97.14 | 35 | 43660 | 8.00E-07 | 58.4 | Paramecium caudatum mitochondrion, complete genome |
| m | scaffold_60762:583 bp | 100 | 31 | 42448 | 7.00E-07 | 58.4 | Coprinopsis cinerea okayama7#130 cont3.68, whole genome shotgun sequence |
| p | scaffold_5546:1007 bp | 100 | 30 | 62891 | 4.00E-06 | 56.5 | Phelipanche purpurea chloroplast complete genome, specimen voucher BONN:S. Wicke Op38/39 |
| m | scaffold_22387:753 bp | 100 | 30 | 704100 | 3.00E-06 | 56.5 | Tripsacum dactyloides mitochondrion, complete genome |
| m | scaffold_134222:448 bp | 100 | 30 | 62978 | 2.00E-06 | 56.5 | Fusarium solani mitochondrion, complete genome |
| m | scaffold_6410:981 bp | 100 | 29 | 978846 | 1.00E-05 | 54.7 | Welwitschia mirabilis mitochondrion, complete genome |
| m | scaffold_20195:771 bp | 100 | 29 | 16975 | 1.00E-05 | 54.7 | Jacana jacana voucher STRI:BC4055 mitochondrion, complete genome |
| p | scaffold_30473:699 bp | 100 | 29 | 147896 | 1.00E-05 | 54.7 | Silene conoidea chloroplast, complete genome |
| p | scaffold_45799:630 bp | 100 | 29 | 165372 | 9.00E-06 | 54.7 | Zygnema circumcarinatum chloroplast, complete genome |
| p | scaffold_47242:625 bp | 100 | 29 | 193197 | 9.00E-06 | 54.7 | Chlorotetraedron incus strain SAG 43.81 chloroplast, complete genome |
| m | scaffold_51060:612 bp | 100 | 29 | 70578 | 9.00E-06 | 54.7 | Saccharomyces pastorianus Weihenstephan 34/70 mitochondrion, complete genome |
| m | scaffold_116963:472 bp | 96.97 | 33 | 30782 | 7.00E-06 | 54.7 | Saccharomyces servazzii mitochondrion, complete genome |
| m | scaffold_141505:439 bp | 92.31 | 39 | 49539 | 6.00E-06 | 54.7 | Rhynchosporium orthosporum mitochondrion, complete genome |
| m | scaffold_209314:365 bp | 100 | 29 | 364070 | 5.00E-06 | 54.7 | Psilotum nudum isolate v16 chromosome 1 mitochondrion, complete sequence |
| m | scaffold_209583:365 bp | 100 | 29 | 16120 | 5.00E-06 | 54.7 | Pyrophorus divergens mitochondrion, complete genome |
| m | scaffold_305995:250 bp | 100 | 29 | 201763 | 3.00E-06 | 54.7 | Chlorokybus atmophyticus mitochondrion, complete genome |
| m | scaffold_5538:1008 bp | 100 | 28 | 53262 | 5.00E-05 | 52.8 | Ogataea thermophila strain NCAIM Y.01608 mitochondrion, complete genome |
| p | scaffold_6935:966 bp | 100 | 28 | 102657 | 5.00E-05 | 52.8 | Cistanche deserticola chloroplast, complete genome |
| m | scaffold_17488:1269 bp | 100 | 28 | 16767 | 7.00E-05 | 52.8 | Castor canadensis mitochondrion, complete genome |
| m | scaffold_20021:772 bp | 100 | 28 | 42781 | 4.00E-05 | 52.8 | Tetradesmus obliquus strain KS3-2 mitochondrion, complete genome |
| p | scaffold_32121:690 bp | 100 | 28 | 145303 | 4.00E-05 | 52.8 | Isoetes flaccida chloroplast, complete genome |
| m | scaffold_49416:618 bp | 100 | 28 | 71124 | 3.00E-05 | 52.8 | Saccharomyces arboricola strain CBS 10644 mitochondrion, complete genome |
| m | scaffold_92258:513 bp | 100 | 28 | 71335 | 3.00E-05 | 52.8 | Paracoccidioides brasiliensis mitochondrion, complete genome |
| m | scaffold_93459:511 bp | 100 | 28 | 982833 | 3.00E-05 | 52.8 | Cucurbita pepo mitochondrion, complete genome |
| m | scaffold_119425:469 bp | 100 | 28 | 31825 | 2.00E-05 | 52.8 | Phakopsora pachyrhizi mitochondrion, complete genome |
| m | scaffold_153656:424 bp | 100 | 28 | 680603 | 2.00E-05 | 52.8 | Zea mays subsp. parviglumis mitochondrion, complete genome |
| m | scaffold_226398:350 bp | 100 | 28 | 105383 | 2.00E-05 | 52.8 | Gaeumannomyces graminis var. tritici R3-111a-1 mitochondrial scaffold supercont1.514, whole genome shotgun sequence |
| p | scaffold_239857:338 bp | 100 | 28 | 171634 | 2.00E-05 | 52.8 | Ceramium japonicum plastid, complete genome |
| p | scaffold_272582:311 bp | 94.29 | 35 | 91517 | 2.00E-05 | 52.8 | Orobanche rapum-genistae chloroplast, complete genome |
| m | scaffold_298848:268 bp | 100 | 28 | 76453 | 1.00E-05 | 52.8 | Dekkera bruxellensis mitochondrion, complete genome |
| m | scaffold_330255:204 bp | 100 | 28 | 51679 | 1.00E-05 | 52.8 | Lachancea kluyveri mitochondrion, complete genome |

**Supplementary Table 4**: Filtered mapped read count for all sequenced *S. pugilis* specimens

| *Individual* | *Site Age* | *Source Material* | *Number of reads mapped* | | | |
|---|---|---|---|---|---|---|
| | | | *Nuclear reads - bwa mem* | *Nuclear reads - bwa aln* | *Mitochondrial reads - bwa mem* | *Mitochondrial reads - bwa aln* |
| BocaDrago3-10 | Modern | Tissue | 10,596,655 | 1,152,019 | 10,751 | 683 |
| BocaDrago3-3 | Modern | Tissue | 6,608,838 | 632,340 | 4,555 | 2,151 |
| CayoAgua2-3 | Modern | Tissue | 51,560,307 | 9,218,590 | 28,023 | 10,970 |
| CayoAgua2-5 | Modern | Tissue | 3,499,281 | 459,756 | 4,329 | 1,315 |
| CayoAgua2-6 | Modern | Tissue | 7,153,558 | 762,193 | 9,389 | 1,826 |
| BocaDrago3-10 | Modern | Shell | 25,361 | 1,697 | 5 | 2 |
| BocaDrago3-3 | Modern | Shell | 147,458 | 8,172 | 10 | 5 |
| CayoAgua2-3 | Modern | Shell | 466,588 | 92,528 | 570 | 192 |
| CayoAgua2-5 | Modern | Shell | 281,446 | 40,506 | 208 | 83 |
| CayoAgua2-6 | Modern | Shell | 218,071 | 24,145 | 51 | 18 |
| CayoAguaBoil1 | Modern | Shell | 1,240 | 525 | 4 | 0 |
| CayoAguaBoil2 | Modern | Shell | 2,488 | 1,063 | 2 | 4 |
| CayoAguaBoil3 | Modern | Shell | 7,647 | 4,574 | 6 | 11 |
| U60-10_20 | Archaeological | Shell | 390 | 3,269 | 0 | 7 |
| U60-110-120-5659 | Archaeological | Shell | 1,138 | 12,060 | 1 | 10 |
| U61-20-30 | Archaeological | Shell | 349 | 1,392 | 0 | 3 |
| U61-50_60-5454 | Archaeological | Shell | 266 | 1,375 | 0 | 2 |
| U61-80_90_5596C | Archaeological | Shell | 476 | 4,459 | 0 | 11 |
| MS-F-10-110 | Paleontological | Shell | 236 | 4 | 0 | 0 |
| MS-F-15-4 | Paleontological | Shell | 195 | 41 | 0 | 0 |
| MS-F-2-78 | Paleontological | Shell | 285 | 9 | 0 | 0 |
| MS-F-5-157 | Paleontological | Shell | 282 | 114 | 0 | 0 |
| MS-F-7-128 | Paleontological | Shell | 293 | 52 | 0 | 0 |

**Supplementary Figure 1**: Nuclear DNA damage and read length results for individual shell samples

### Individual Nuclear DNA Damage and Read Lengths

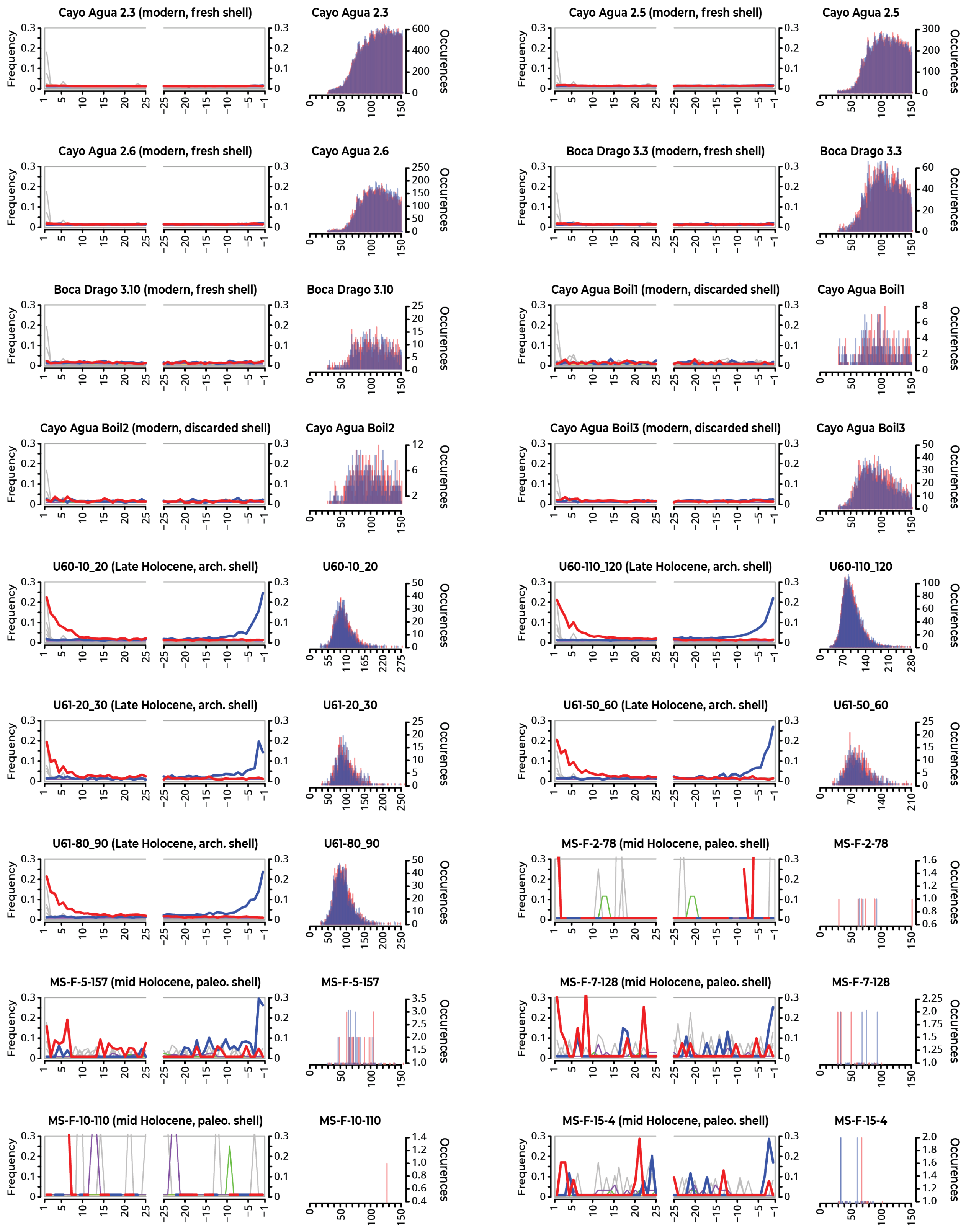
